# Coupling and uncoupling of midline morphogenesis and cell flow in amniote gastrulation

**DOI:** 10.1101/2023.05.26.542486

**Authors:** Rieko Asai, Vivek N. Prakash, Shubham Sinha, Manu Prakash, Takashi Mikawa

## Abstract

Large-scale cell flow characterizes gastrulation in animal development. In amniote gastrulation, particularly in avian gastrula, a bilateral vortex-like counter-rotating cell flow, called ‘polonaise movements’, appears along the midline. Here, through experimental manipulations, we addressed relationships between the polonaise movements and morphogenesis of the primitive streak, the earliest midline structure in amniotes. Suppression of the Wnt/planar cell polarity (PCP) signaling pathway maintains the polonaise movements along a deformed primitive streak. Mitotic arrest leads to diminished extension and development of the primitive streak and maintains the early phase of the polonaise movements. Ectopically induced Vg1, an axis-inducing morphogen, generates the polonaise movements, aligned to the induced midline, but disturbs the stereotypical cell flow pattern at the authentic midline. Despite the altered cell flow, induction and extension of the primitive streak are preserved along both authentic and induced midlines. Finally, we show that ectopic axis-inducing morphogen, Vg1, is capable of initiating the polonaise movements without concomitant PS extension under mitotic arrest conditions. These results are consistent with a model wherein primitive streak morphogenesis is required for the maintenance of the polonaise movements, but the polonaise movements are not necessarily responsible for primitive streak morphogenesis. Our data describe a previously undefined relationship between the large-scale cell flow and midline morphogenesis in gastrulation.

## Introduction

Large-scale cell flow during gastrulation is an evolutionarily conserved biological phenomenon in embryogenesis (1–3). During this period, animals with bilateral symmetry, called bilaterians, initiate development of three germ layers (i.e. the ectoderm, mesoderm and endoderm) and midline structures (e.g. the notochord and primitive streak) along the midline axis (4–6). The large-scale cell flow has been linked to early embryonic morphogenesis, including axial structures, through rearrangement of cells and/or transportation of signaling molecules (2, 3). Coupling of the large-scale cell flow and body axis morphogenesis has been extensively studied in invertebrate bilaterians, particularly insect models (7–9). However, in vertebrates, our understanding of whether and how the large-scale cell flow and midline morphogenesis are coupled is limited.

Vertebrates are classified into two groups: non-amniotes (e.g. fish and amphibians) and amniotes (e.g. birds and mammals) (1). In non-amniotes, particularly amphibians, the large-scale cell flow is coupled with formation of the notochord through Wnt/planar cell polarity (PCP) pathway-regulated convergent extension, which does not depend on mitosis (10, 11). In amniotes, particularly in avian gastrula (i.e. embryonic disc), a bilateral vortex-like counter-rotating cell flow, termed ‘polonaise movements’, occurs within the epiblast along the midline axis, prior to and during primitive streak (PS) formation (12–14). The PS is an evolutionarily unique embryonic structure in amniotes, which is not only an organizing center for gastrulation but also the earliest midline structure (6, 15–17). During amniote gastrulation, widespread mitosis has been identified throughout the embryo, including the PS (18), and previous studies showed that mitotic arrest leads to morphologically diminishing PS extension (19, 20). Further, the Wnt/PCP pathway also plays an important role for proper PS extension and patterning (21). Since the chick embryo is readily accessible and closely represents the human gastrula (22, 23), much of what is known about the cell flow and PS development has come from studies using this model system (14). The PS is first visible as a cell cluster at the posterior embryonic disc at the initial streak stage [HH2; Hamburger and Hamilton staging (24)] formed through the influence of several axis inducing factors [e.g. Vg1 and Chordin (25, 26)]; it then extends anteriorly along the midline axis (6, 14, 15). During PS development, displacement of cells and/or tissue tension potentially generates biomechanical forces, which lead to cell flows and/or PS morphogenesis (20, 21, 27, 28). Recent studies discuss that cell intercalation and ingression underlying PS formation, leading to the polonaise movements through altering the neighbor tissue tension (21, 28, 29), suggesting that the polonaise movements are likely a consequence of PS morphogenesis (16, 30, 31). Other studies demonstrate that the polonaise movements, generated by the myosin-cable-mediated tissue tension with cell intercalation and ingression, lead to PS morphogenesis (20, 27, 31). Thus, the relationship between the polonaise movements and PS morphogenesis remains controversial (31).

Here, we characterize previously undefined relationships between the polonaise movements and PS morphogenesis. Suppression of the Wnt/PCP pathway generates a wider and shorter PS while maintaining the polonaise movements. Mitotic arrest diminishes PS extension and maintains the early phase of the polonaise movements. Moreover, experimentally manipulating the large-scale cell flow by inducing a secondary midline axis indicates that the authentic PS forms despite altered cell flows. Ultimately, a single axis-inducing morphogen, Vg1, is capable of initiating the polonaise movements despite defective PS morphogenesis. These results suggest that PS morphogenesis is not responsible for the initiation and early phase of the polonaise movements. Further, the polonaise movements are not necessarily responsible for PS morphogenesis. This study suggests that amniote gastrulation may be evolutionarily distinct from non-amniote gastrulation in the mechanisms coupling large-scale cell flow and midline morphogenesis.

## Results

### Suppression of the Wnt/PCP pathway maintains the bilateral vortex-like counter-rotating cell flow along a deformed PS

Throughout this work, the cell flows were recorded by tagging individual epiblast cells with electroporated fluorescent tracers at the pre-streak stage (32). The tagged embryos were live-imaged until HH3 with a modified New culture system, which we previously described (33, 34). To reconstruct trajectories of the tagged cells and quantitatively analyze the cell flow pattern, the recorded images were processed using tracking analysis tools (35), Particle Image Velocimetry (PIV) (36–40), and subsequent visualization techniques (41, 42).

To address the relationship between the polonaise movements and PS morphogenesis, we experimentally manipulated the shape of the PS while maintaining the population of PS cells (Fig. 1A-B). The DEP domain of Dishevelled (Dsh; a transducer protein of Wnt signaling) is responsible for the non-canonical Wnt/PCP pathway (43, 44), and misexpression of dominant-negative Dsh lacking DEP [dnDsh(ΔDEP)] leads to deformation of the midline structures, including the PS (21). Further, the Wnt/PCP pathway is involved in cellular polarity and migration, while the canonical Wnt pathway regulates cell proliferation (45). We refer the dnDsh(ΔDEP)-GFP construct that we generated, as ΔDEP-GFP, and tested its ability to alter cellular polarity, resulting in PS deformation’ (Fig. 1C-E, Fig. S1A-D). In the control-GFP-introduced PS and epiblast cells, Pk1 (a PCP core component) exhibited a polarized localization at the E-cadherin-based adherens junctions, particularly cell-cell junctions (Fig. S1A-B, n=3). In contrast, this Pk1-localization pattern was not identified in the ΔDEP-GFP-misexpressing PS and epiblast cells (Fig. S1C-D, n=3), implying that the construct efficiently disrupted the cellular polarity within the misexpressing cells. The expression of *Brachyury* indicated that ΔDEP-GFP-PS cells maintained their identity (Fig. 1E) but displayed a shorter and wider morphogenesis than the control PS (Fig. 1C-E, n≥4 for each). The number of PS cells was not significantly different from the control- and ΔDEP-GFP-PS (control 8.5±0.4×10^3^ cells vs ΔDEP 8.8±1.2×10^3^ cells, p=0.68, n=3 for each, Fig. 1F). These results indicated that ΔDEP-misexpression was capable of suppressing the Wnt/PCP pathway through disrupting the localization pattern of the PCP core component while maintaining the number of the PS cells. Having confirmed the function of the ΔDEP-GFP construct, we applied it as a molecular tool for manipulating the shape of the PS.

**Fig. 1.**
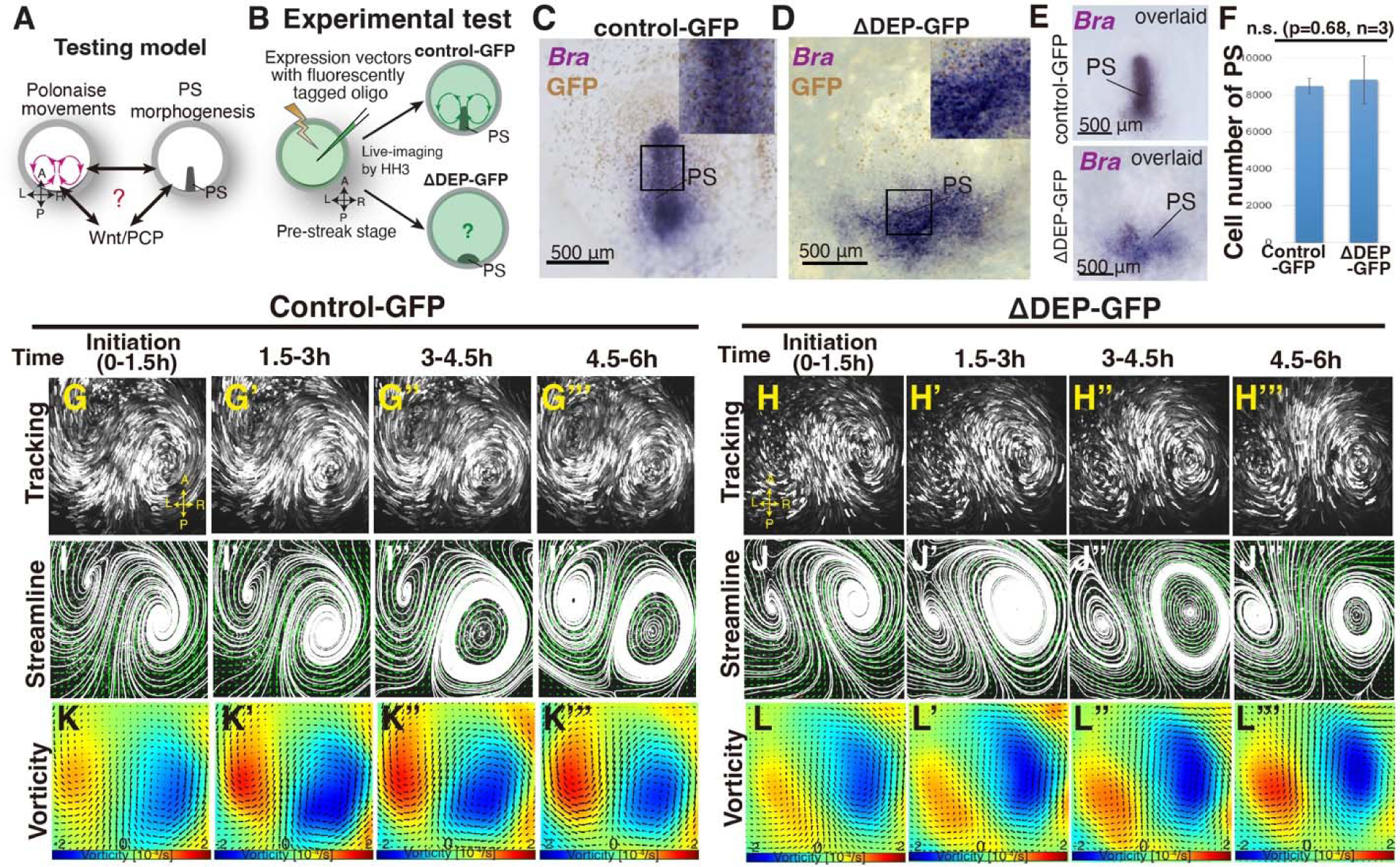
‘Polonaise movements’ persist under suppression of Wnt/PCP pathway. (A) Testing model of relationships between the polonaise movements, primitive streak (PS) morphogenesis, and the Wnt/PCP pathway. A-P and L-R; anterior-posterior and left-right body axes, respectively. (B) Diagram of experimental set up. (C, D) In situ hybridizations for Brachyury (Bra) in control- and ΔDEP-GFP-misexpressing embryos. (E) Overlay of embryos hybridized for Bra (n=4 for each). (F) Cell number in PS [control 8.5±0.4×10^3^ cells vs ΔDEP 8.8±1.2×10^3^ cells, p=0.68, n=3 for each (two-tailed Student’s t test)]. (G-H’’’) Trace of cell flow path of electroplated epiblast cells with Flowtrace. The initiation of the polonaise movements was set as Time 0 (t=0). See also Movie S1. (I-J’’’) Streamlines, visualizing averaged cell flows during each time period. (K-L’’’) Vorticity plots, displaying an averaged measure of the local rotation during each time period. Blue, clockwise; red, counter-clockwise rotation.

To address relationship(s) between the polonaise movements and the shape of the PS, we live-imaged the control- and ΔDEP-GFP-embryos from pre-streak stage to HH3 and analyzed the resulting cell flows (Fig. 1G-L”’, and movie S1). In the control-GFP embryo, the cell flow pattern, which was visualized by tracking tools and quantitatively analyzed by PIV with subsequent flow-visualization techniques (41, 42), identified robust polonaise movements (Fig. 1G-G”’, I-I”’, K-K”’, and movie S1, n=5). Surprisingly, these analyses indicated that suppression of the Wnt/PCP pathway maintained the polonaise movements despite PS deformation (n=5, Fig. 1C-L”’ and movie S1). While the topology of the counter-rotating flow pattern were maintained under suppression of the Wnt/PCP pathway (Fig. 1J-J’”), the distance between the left-right rotating cell flows was wider in the ΔDEP embryos than the controls (control 860.2±86.2μm vs ΔDEP 1008.4±183.8μm, p=0.026, n=4 for each, Fig. S1E-I), implying that the shape of the PS and/or the Wnt/PCP signaling-mediated cell intercalation affected the location of these rotations. These data demonstrated that the polonaise movements persisted even though the PS as a midline structure was deformed under the condition of suppressing the Wnt/PCP pathway.

### Mitotic arrest diminishes PS formation but initially preserves the bilateral vortex-like rotating cell flow

The above data showed that a deformed midline structure allowed the polonaise movements to occur while the position of the LR rotations in the polonaise movements was affected (Fig. 1 and Fig. S1). However, it remained unclear the relationship between the polonaise movements and the population of the PS cells, particularly expansion of the PS precursor cells in PS morphogenesis (Fig. 2A-B). To address this question, we applied a mitotic arrest approach using aphidicolin (a specific DNA polymerase inhibitor), which has been shown to diminished PS development (19, 20). In chick embryonic disc, active cell proliferation was identified in both the epiblast and PS cells (Fig. 2C)(18). As verified by BrdU incorporation assay, the aphidicolin-treated embryos exhibited little or no mitotic activity (Fig. 2D). Morphogenesis of the PS was monitored by *Brachyury* (a marker for PS and mesodermal cells) and *Sox3* (an epiblast marker). The expression domain and midline extension of cells, expressing *Brachyury* and negative for *Sox3*, were considerably reduced in the aphidicolin-treated embryos, compared to the control (Fig. 2E; n≥4 for each). The expression pattern of *Brachyury* and PS extension followed an aphidicolin dose-dependent diminishment (Fig. S2-D). The *Vg1*-expressing area was also diminished but detectable in the aphidicolin-treated embryos (Fig. 2E, n=4), suggesting that the axis-inducing morphogen was preserved in the embryonic disc, but expansion of the PS precursor cells was suppressed by mitotic arrest. These results indicate that mitotic arrest, induced by the aphidicolin-treatment, remarkably reduced PS extension and development, consistent with the previous works (19, 20).

**Fig. 2.**
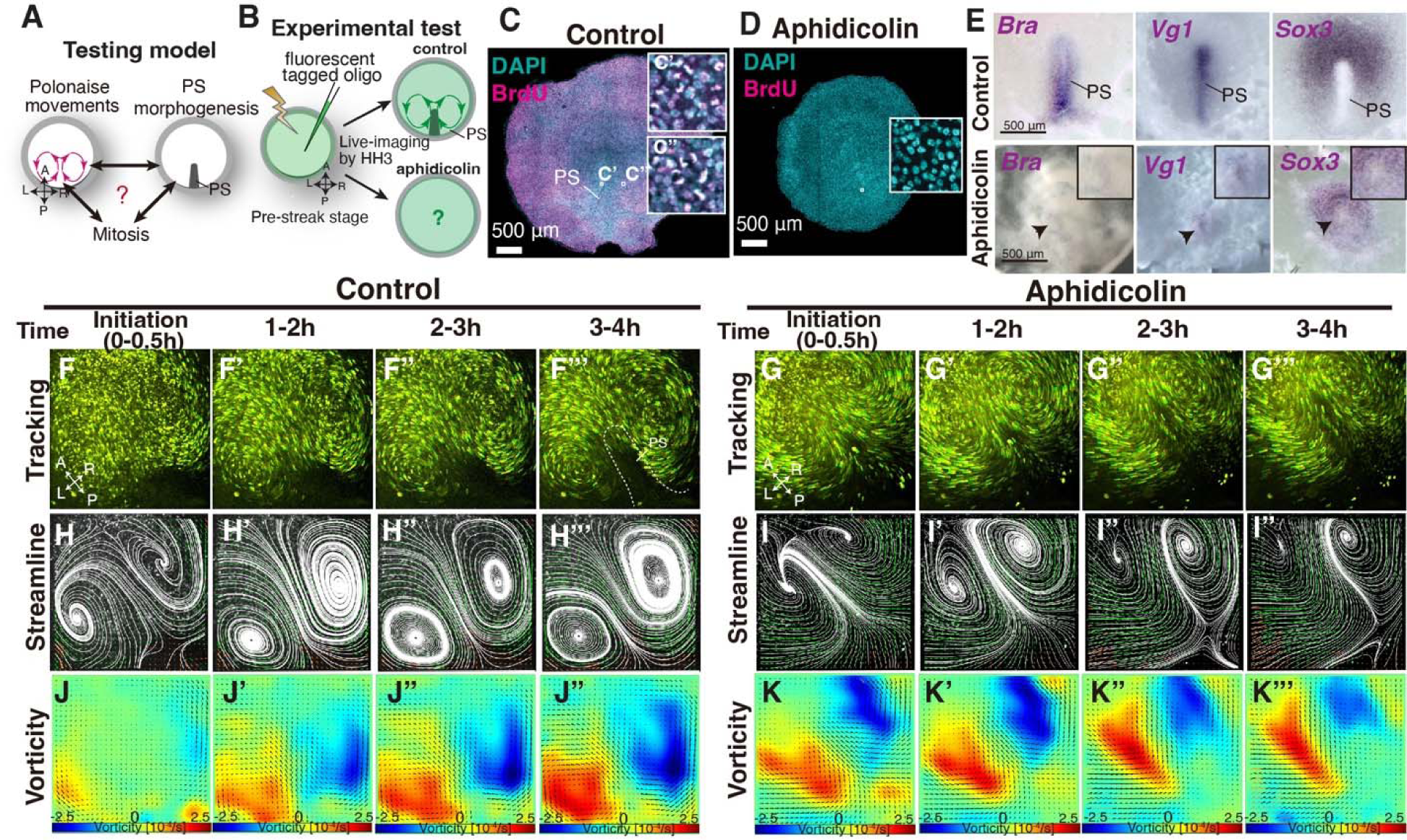
Early phase of ‘polonaise movements’ remains under global mitotic arrest. (A) Testing model of relationships between the polonaise movements, primitive streak (PS) morphogenesis, and mitosis. Axes as in Fig. 1. (B) Diagram of experimental set up. (C, D) BrdU incorporation in control-sham operated and aphidicolin treated in embryos. White boxes show enlarged areas. (C’, C”) High magnification of boxed areas in PS and non-PS, respectively. (E) In situ hybridizations for *Brachyury* (*Bra*), *Vg1*, and *Sox3* in control and aphidicolin-treated embryos. Arrow heads indicate diminished PSs. Black boxes, enlarged area including the diminished PS. (F-G’’’) Trace of cell flow path with Flowtrace analysis (green is trace, yellow indicates endpoint) of fluorescently-tagged epiblast cells. The initiation of the polonaise movements was set as Time 0 (t=0). See also Movie S2. (H-I’’’) Streamlines, visualizing averaged cell flows during each time period. Interpolated vectors are displayed in orange. (J-K’’’) Vorticity plots, displaying averaged measure of the local rotation during each time period. Blue, clockwise; red, counter-clockwise rotation. **Note**: Since vorticity is calculated for all deviation from set point, slight curves and full rotations receive the same color indication, as seen in K’’’ which is not maintaining a bilateral vortex-like-rotating cell flows. See Materials and Methods section for a full discussion.

To address the relationship between the polonaise movements and the population of the PS cells, we live-imaged the fluorescently tagged embryos under aphidicolin-treatment and analyzed the resulting cell flow (Fig. 2F-K”’ and movie S2). Tracking and PIV-based analyses visualized the cell flow pattern in control and aphidicolin-treated embryos (Fig. 2F-K”’ and movie S2). In control embryos, the polonaise movements occurred from pre-streak until HH3 for more than four hours (n=4, Fig. 2F-F”’ and movie S2). The streamline and vorticity plots illustrated that the polonaise movements became more robust in a time-dependent manner (Fig. 2H-H”’ and J-J”’). Strikingly, in the aphidicolin-treated embryos (n=4), the polonaise movements initiated but were maintained for a shorter period, 2.2±1.2 hours (control 6.5±1.2 hours, p=0.0038, Fig. G-G’”, I-I’”, and movie S2). The streamline and vorticity plots also indicated the initiation and early phase of the polonaise movements in the aphidicolin-treated embryos (Fig. 2J-J”’ and K-K’”). Together, these data demonstrated that the typical pattern of the polonaise movements was induced and initially maintained in the short-term, while there was no or little indication of robust PS formation or extension.

Aphidicolin potentially induces apoptosis (46), which might indirectly lead to the cell flow changes through tissue tension alterations. To test this possibility, we examined apoptosis in the control and aphidicolin-treated embryos (Fig. S3A-B). TUNEL assay showed that the aphidicolin-treatment slightly increased apoptotic rate but there was no significant difference between these embryos and controls (control sham, 10.6±3.5%; aphidicolin, 12.1±3.7%, p=0.20, n=4 for each, Fig. S3B), suggesting that the bilateral rotating cell flow in the aphidicolin-treated embryos was not indirectly induced by apoptosis.

Previous studies have shown that a large-scale ring-like distribution of phospholyrated-myosin (pMyosin) cables among the epiblast cell layer is required for tissue tension related to cell flow (20, 27, 28). We examined whether the pMyosin cables were maintained in the treated embryos over the course of the imaging (Fig. S3C-D”). In the aphidicolin-treated embryos, the large-scale ring-like distribution pattern of the pMyosin cables remained in the epiblast (Fig. S3D-D”), similar to the control embryos (Fig. S3C-C”), implying that the mitotic arrested-embryos maintained tissue tension, consistent with the previous report (20). Thus, these results support a model that mitosis is coupled with PS morphogenesis but is not necessarily required for the initiation and early phase of the polonaise movements.

### PS properly extends along the midline despite altering the large-scale cell flow

Given that PS morphogenesis is required for maintenance of the polonaise movements, we next examined the converse of the relationship; that the polonaise movements may be required for PS formation. To test this model, we experimentally reprogrammed the large-scale cell flow (Fig. 3A-B). Ectopically induced Vg1 leads to formation of a *Brachyury*-positive secondary PS and a secondary midline axis (Fig. 3C-D)(47–49). COS cells expressing either a control-GFP (control-GFP/COS) or Vg1 (Vg1/COS) construct were placed at the anterior marginal zone (AMZ) at pre-streak stage, approximately 180° from the posterior marginal zone where the authentic PS forms, and we analyzed the resulting cell flow during both authentic and ectopically induced PS extension (Fig. 3B). If the bilateral rotating cell flow is required for PS formation, both the authentic and induced PSs should each be accompanied by polonaise movements during the extension. The trajectory analysis by Imaris and PIV-based analyses showed that the control-COS implanted embryos exhibited stereotypical polonaise movements along the authentic midline axis, indicating that the COS cell implantation maintained an authentic cell flow pattern (Fig. 3E-E’”, G-G’”, I-I’”, K, Fig. S4, and movie S3, n=4). In contrast, the analyses of the cell flow in the Vg1/COS implanted embryos illustrated the disruption of the authentic polonaise movements at the extending authentic PS (Fig. 3F-F”’, H-H”’, J-J”’, Fig. S5, and movie S3, n=4). At pre-streak stage, the epiblast cells initially converged to the center of the embryonic disc along both the authentic and the induced axes, and the bilateral rotating cell flows began to appear at both axes (Fig. 3F-F’”, H-H’”, Fig. S5, and movie S3). This cell flow pattern continued for approximately one hour (Fig. 3H’, Fig. S5, and movie S3). Before extension of the authentic PS, the bilateral rotating cell flow along the induced axis became dominant, and eventually overrode the original cell flow (Fig. 3H’-H’”, Fig. S5, and movie S3). Despite global disruption and reversed counter-rotating cell flows, the authentic PS extended properly along the authentic midline (Fig. 3D, F”’, H”’, J”’, K, Fig. S5, and movie S3; n=4). Secondary midline-axis induction by ectopic Vg1 created the polonaise movements around the secondary PS and altered the original cell flow pattern, generating a reprogrammed large-scale cell flow (Fig. 3F-F”’, H-H”’, J-J”’, K, Fig. S6, and movie S3). These results indicate that extension of the authentic PS is preserved even under the defective polonaise movements.

**Fig. 3.**
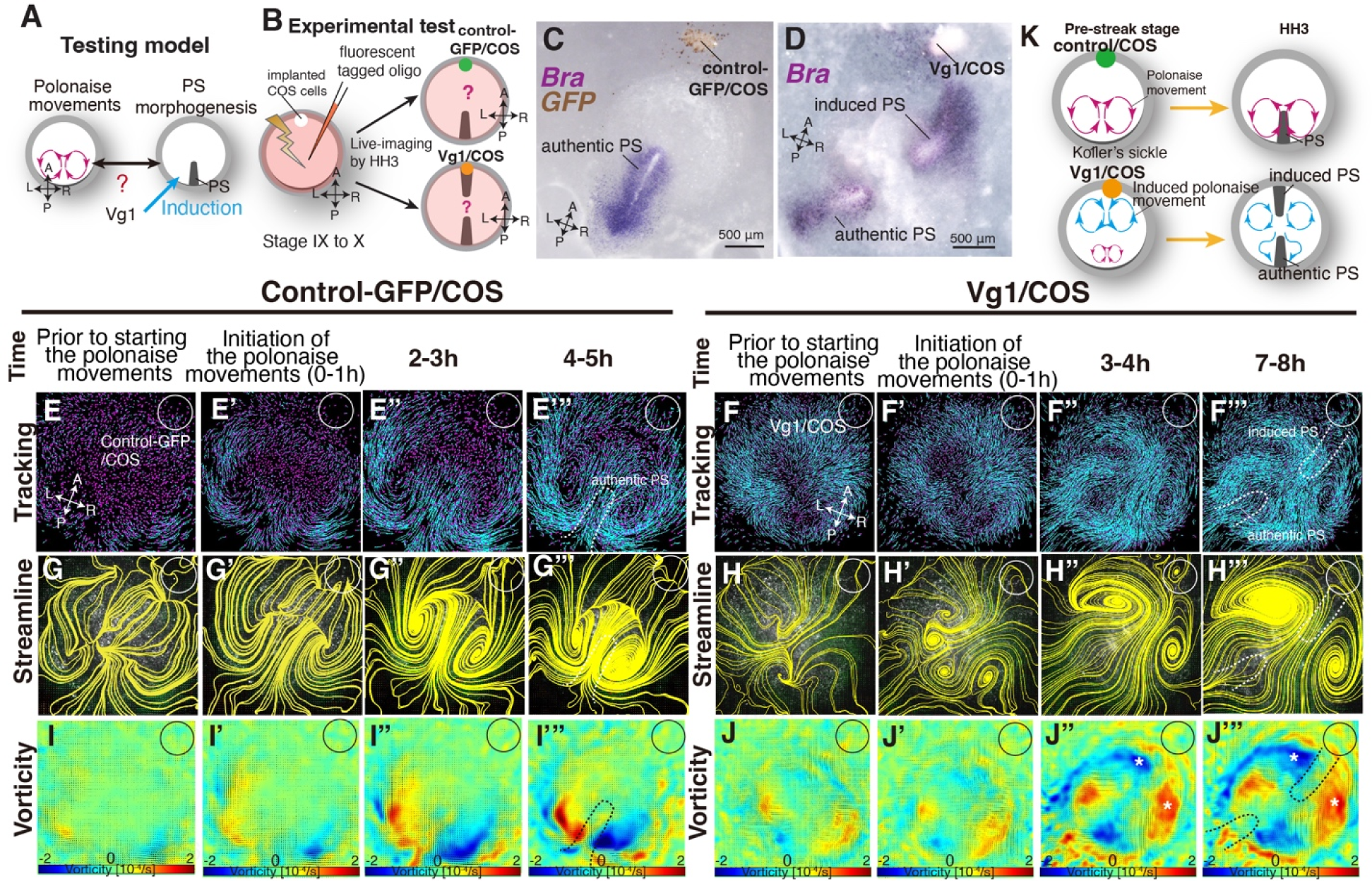
Authentic PS extends under disruption of authentic ‘polonaise movements’. (A) Testing model of relationships between the polonaise movements, primitive streak (PS) morphogenesis, using ectopic Vg1-inducing system. Axes as in Fig. 1. (B) Diagram of experimental set up. (C, D) In situ hybridizations for Brachyury (Bra) in control and ectopic Vg1-induced PS, respectively. (E-F’’’) Trace of cell flow path of electroplated epiblast cells by using Imaris tracking analysis. See also Movie S3. White circles, COS cell implanted site. The initiation of the polonaise movements was set as Time 0 (t=0). (G-H’’’) Streamlines, visualizing averaged cell flows during each time period of the vector fields. (I-J’’’) Vorticity plots, displaying an averaged measure of the local rotation during each time period in the vector fields. Blue, clockwise; red, counter-clockwise rotation. **Note**: vorticity is identified in both a full-rotation and curve, particularly in (J-J’’’). The white asterisks indicate the polonaise movements at the induced PS which are opposite direction to the authentic PS. (K) Summary of the cell flow patterns in control-GFP/COS- and Vg1/COS-implanted embryos, shown in (E-J’”).

### Vg1 is capable of inducing the bilateral vortex-like rotating cell flow despite diminished PS extension under mitotic arrest

The above data indicate the lack of a strong relationship between the polonaise movements and PS morphogenesis; however, it remained unclear how the polonaise movements were induced. Given that mitotic arrest maintained both Vg1-expression and the initiation of the polonaise movements (Fig. 2 and movie S2) and that induction of the secondary midline axis by ectopic Vg1 reprograms the epiblast cell flow (Fig. 3, Fig. S4, S5, and movie S3), we tested the possibility that the axis-inducing morphogen, Vg1, would be capable of inducing the polonaise movements in the absence of PS extension (Fig. 4A). We combined the secondary axis induction by AMZ implantation of Vg1/COS cells with aphidicolin-treatment and analyzed the resulting cell flow (Fig. 4B, Fig. S6, S7, and movie S4). The BrdU assay showed that the aphidicolin treatment effectively blocked mitosis in Vg1/COS implanted embryos (control sham 88±9.0% vs control-GFP/COS with aphidicolin 0.2±0.1% vs Vg1/COS with aphidicolin 0.19±0.2%, p= 0.0005×10^-3^, n=3 for each, Fig. 4C-D) and diminished both the authentic and induced PSs (Fig. 4E-F, n=3 for each). The control-GFP/COS implanted embryo under mitotic arrest condition initially displayed the polonaise movements along the authentic midline axis, however, the vorticity decreased time-dependently, resulting in a defective rotating cell flow (n=3, Fig. 4G-G”’, I-I”’, K-K”’, Fig. S6, and movie S4), consistent with the non-COS cell implanted embryos under aphidicolin-treatment (shown in Fig. 2). In contrast, the Vg1/COS implanted embryos under mitotic arrest condition exhibited the polonaise movements predominantly at the induction point of the secondary axis with disrupted authentic cell flow at the site of the authentic midline axis (n=3, Fig. 4H-H”’, J-J”’, L-L”’, and movie S4). These data suggest that the midline axis induction by the single morphogen brings about the polonaise movements, even when the midline structure (i.e. extending PS) is diminished. Taken together, our data show that Vg1 is capable of inducing and reprogramming the large-scale cell flow during gastrulation even though the midline structure is diminished.

**Fig. 4.**
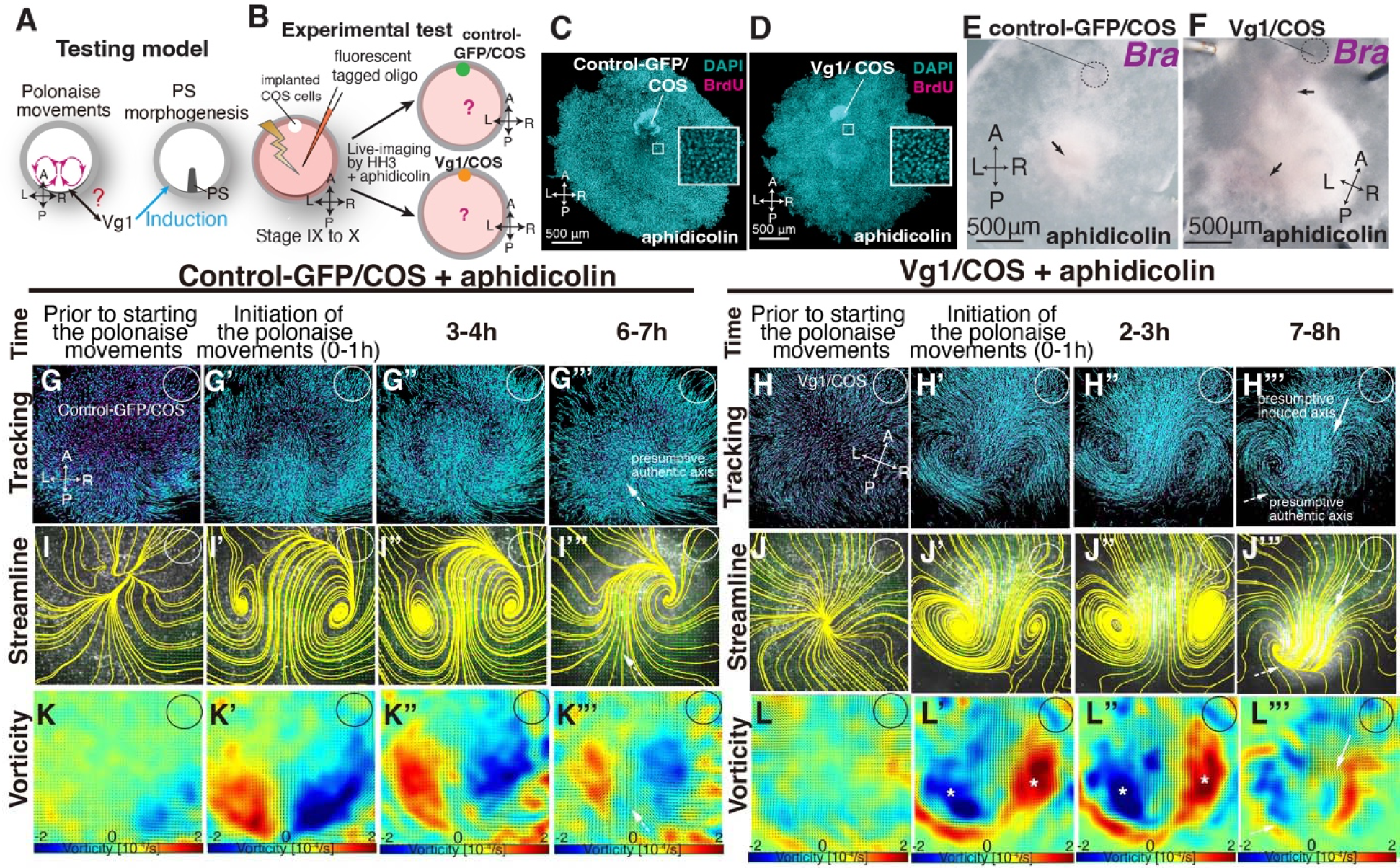
Induced ‘polonaise movements’ persist despite defective PS morphogenesis. (A) Testing model of relationships between the polonaise movements, primitive streak (PS) morphogenesis, and Vg1. Axes as in Fig. 1. (B) Diagram of experimental set up. (C, D) BrdU incorporation in control-GFP/COS- and Vg1/COS-implanted embryos under aphidicolin-treatment. White boxes showed enlarged areas. (E, F) In situ hybridizations for Brachyury (Bra, n=3 for each). Black arrows indicate diminished PSs. (G-H’’’) Trace of cell flow path of electroplated epiblast cells by using Imaris tracking analysis. See also Movie S5. White circles, COS cell implanted site. The initiation of the polonaise movements was set as Time 0 (t=0). (I-J’’’) Streamlines, visualizing averaged cell flows during each time period of the vector fields. (K-L’’’) Vorticity plots, displaying an averaged measure of the local rotation during each time period. Note: vorticity is identified in both a full-rotation and curve, particularly in (L-L’’’). Blue, clockwise; red, counter-clockwise rotation. White asterisks, the polonaise movements at the induced axis which are opposite direction to the authentic midline axis. **Note:** vorticity is identified in both a full-rotation and curve in (K’’’) and (L’’’). See description in the material and method section.

## Discussion

Here, we described a previously undefined relationship between the polonaise movements and PS morphogenesis (Fig. 5). Our approaches to molecularly manipulating PS morphogenesis revealed that the initiation and early phase of the polonaise movements occur despite diminished PS formation (Fig. 5A). Conversely, the experimental manipulation of the cell flow demonstrated that the authentic PS extends regardless of aberrant flow patterns (Fig. 5B). Further, a single axis-inducing morphogen, Vg1, is capable of inducing the bilateral rotating cell flow even under mitotic arrest, when the ectopic PS is reduced (Fig. 5B, C). Our data support a model in which the PS population is required for maintaining the polonaise movements, as seen in the ΔDEP embryos, while the polonaise movements are not necessarily responsible for PS morphogenesis (Fig. 5D).

**Fig. 5.**
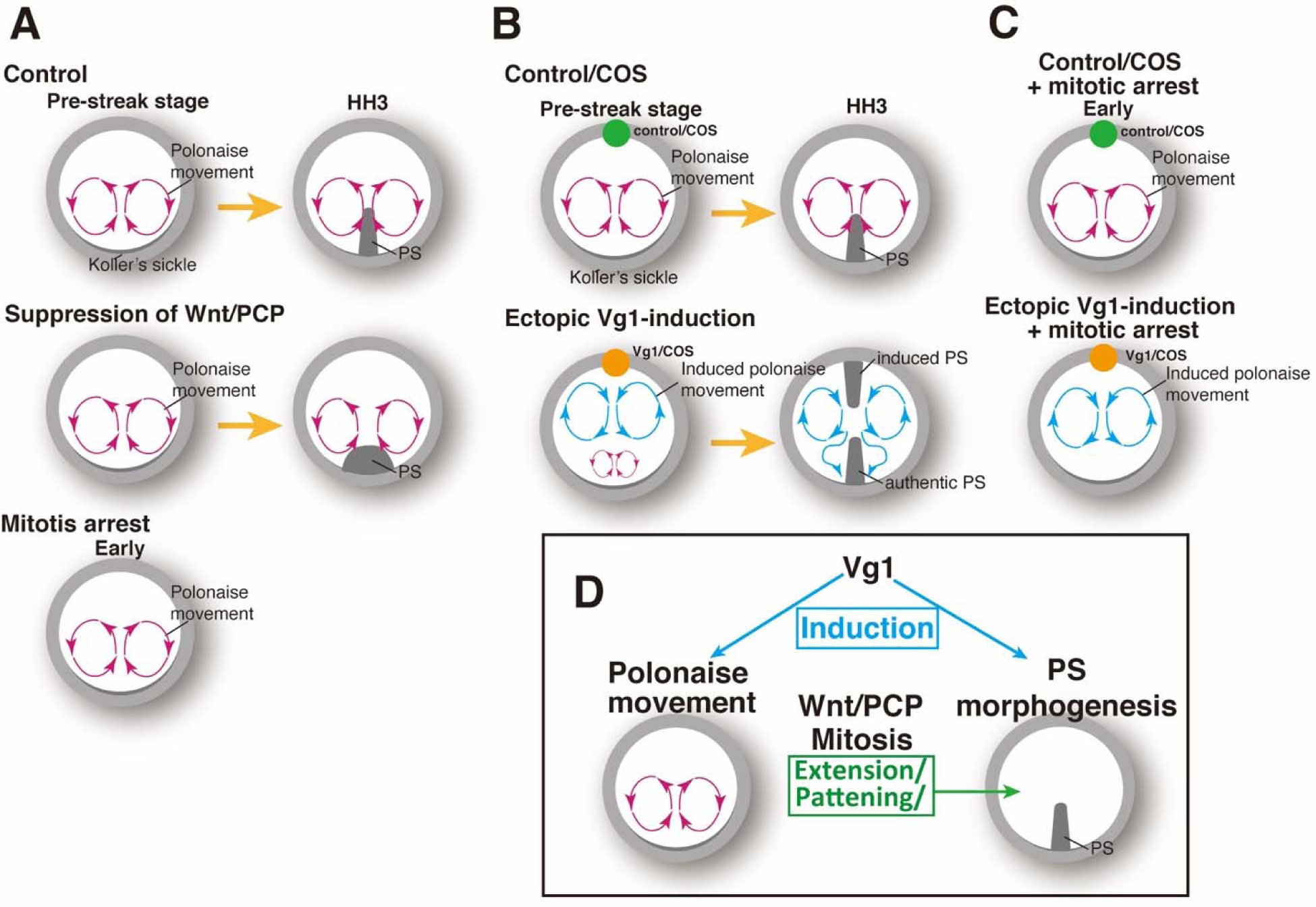
Summary of the patterns of the cell flow and PS morphogenesis. (A) Cell flow pattern under experimental manipulation of PS morphogenesis. (B) Authentic PS morphogenesis under disrupting authentic polonaise movements. (C) Vg1-induced cell flow pattern under defective PS morphogenesis. (D) Summary of the relationships between polonaise movements, PS morphogenesis, mitosis, and Wnt/PCP pathway.

A similar mitotic arrest has been described as simultaneously diminishing PS extension and disrupting the polonaise movements (19, 20), but it was not determined whether a lack of PS extension and/or diminished cell population of the PS at the midline led to defective polonaise movements. In this study, our live-imaging system combined with the quantitative analyses, molecular tools for manipulating PS formation, and the gene expression analyses allowed us to clarify the relationships among PS morphogenesis, the polonaise movements, mitosis, and the Wnt/PCP pathway (Fig. 1, Fig. 2, and Fig. 5A, C). Our data demonstrated that the shape of the PS as a midline structure is not necessarily responsible for initiation of the polonaise movements while the population of the PS cells, expanded by mitosis, is required for sustained polonaise movements (Fig. 1 and 2). Mesendoderm specification is required for large-scale cell flows associated with later stage ingression and intercalation at the PS during gastrulation (27, 50). Here, we have described the distinctive set of cell flows that precede gastrulation in the chick embryo, and the relationship between early and late cell flow has yet to be examined.

The Wnt/PCP pathway has been assumed to be coupled with the polonaise movements, as an evolutionarily conserved mechanism (21, 51). In contrast, another study showed that misexpression of dnWnt11 or dnDsh-ΔPDZ [suppressing both the Wnt canonical and non-canonical pathway, including the Wnt/PCP (43)] maintained the bilateral counter-rotating cell flow along a defective PS (52). Thus, the role of the Wnt/PCP signaling pathway for the polonaise movements has remained unresolved. In this study, our loss-of-function technique, introducing ΔDEP to PS cells, demonstrated that the polonaise movements were preserved along the deformed PS (Fig. 1, and movie S1). Our electroporation-mediated gene-transfection allowed us to restrict gene transduction in area and number of cells (53). Despite the technical limitations, the suppression level of the Wnt/PCP pathway through the ΔDEP-misexpression was sufficient to lead to the PS deformation (Fig. 1 and Fig. S1). The Wnt/PCP pathway potentially works both cell-autonomously and non-cell autonomously (45, 54). In the PS and non-PS epiblast cells, the disruption of the Pk1-localization was identified not only in the ΔDEP-misexpressing cells, but also in neighbors; however, this study is not designed to provide insight into questions of cell autonomy. Further, the Wnt/PCP pathway is a major regulator of cell intercalation (51, 55). Recent studies have shown that cell intercalation plays an important role in PS morphogenesis and the initiation of the polonaise movements (20, 21, 27–29). Our data demonstrates that suppression of the Wnt/PCP pathway preserves the initiation and maintenance of the polonaise movements, but the lateral positioning of the rotations is enlarged when the PS is deformed (Fig. 1 and Fig. S1). These results suggest that the Wnt/PCP pathway may not be associated with directional cell intercalation for driving the polonaise movements (Fig. 5C).

To investigate the role of the polonaise movements for PS morphogenesis, a technical challenge/limitation is how to stop or disrupt the cell flow while preserving the cellular process intact. For example, previous loss-of-function studies, using knockdown techniques (such as siRNA and inhibitor treatments), identified the necessity of myosins, including pMyosin and actomyosin, for both the large-scale cell flow and PS morphogenesis, through regulation of tissue tension/biomechanical force (27, 28). Myosins are fundamental cellular components and involved in several cell biological processes, such as tissue tension, cell division, and intracellular transport (56). Therefore, it remains a possibility that myosin-related mechanisms could be involved in the cell flow and PS morphogenesis. Recent studies using injury and pressure models further demonstrated that the cell flow pattern can be altered by a physical cut to an embryonic disc or removal of a large portion of an embryo disc through change of the tissue tension or ablation (20, 57). While these models suggest that the cell flow pattern is dependent on an intact embryonic architecture, it remains unclear as to involvement of other elements such as signaling pathways and gene expression, which can be also molecularly and cell-biologically changed as a result of surgeries. Thus, we developed an experimentally manipulation of the cell flow system, using a Vg1-induced two-midline axis model, disturbing the stereotypical cell flow pattern while maintaining embryonic development (Fig. 3 and Fig. 5B).

Our two-midline axis model provided an important insight into the axis-inducing morphogen for induction and patterning of the polonaise movements (Fig. 3, Fig. 4, Fig. S4-S7, and movie S3, S4). Our data demonstrated that the ectopic Vg1, mimicking the inducing activity of the posterior marginal zone, generated the bilateral rotating cell flow aligned to the induced midline axis but disrupted the cell flows at the authentic midline axis (Fig. 3 and Fig. 4). Despite the disturbed cell flow, PS extension was preserved along both authentic and induced midline axes (Fig. 4). While this model ultimately disturbed the stereotypical cell flow pattern at the authentic midline axis, cell flow was initiated there. Therefore, our study does not rule out potential roles of the cell flow in the initiating steps of PS extension.

It remains unclear how the induced polonaise movements along with the secondary axis overcame the authentic cell flows despite mitotic arrest (Fig. 4 and Fig. 5). As for axis-induction by other axis-inducing morphogens (58), Vg1 may work dose-dependently for induction and patterning of the polonaise movements, and ectopic application may have over-ridden endogenous levels (Fig. 3 and Fig. 4). Further studies will clarify the relationship between the large-scale cell flows and the concentration of the morphogen; this will add new insight to the regulation mechanisms that connect axis-induction and the cell flow patterning.

This study further suggests evolutional distinctions underlying cell biological mechanisms of the large-scale cell flow and midline morphogenesis, between amniotes and non-amniotes. The first distinction is the necessity of the Wnt/PCP pathway to the large-scale cell flow. Whereas the Wnt/PCP pathway is coupled with both the large-scale cell flow and midline morphogenesis in non-amniotes (55), our data revealed that the signaling pathway is required for proper PS morphogenesis, but contributes little to the polonaise movements (Fig. 1). The second distinction is mitosis-dependency/independency comparisons between model systems of gastrulation (Fig. 2). Non-amniotes, in particular amphibians, utilize a blastopore-mediated gastrulation and undergo notochord formation with the large-scale cell flow, all independent of mitosis (10). In amniotes, mitosis is required for PS morphogenesis, and expanding the mass of the PS cells contributes to the maintenance of the polonaise movements (Fig. 2). Thus, the large-scale cell flow and midline morphogenesis in amniote gastrulation is not tightly coupled, which may be evolutionarily distinct from non-amniote gastrulation.

In summary, our study illustrates the relationships between midline morphogenesis, the large-scale cell flow, and an axis-inducing factor in amniote gastrulation. These findings will add another layer of our understanding to how the bilateral body plan is initiated and patterned along the midline axis.

## Acknowledgments

We thank past and present Mikawa lab and Prakash lab members, particularly Drs J. Hyer and L. Hua for their invaluable suggestions and/or assistance. Imaging data for this study were acquired at the Center for Advanced Light Microscopy-CVRI at UCSF on microscopes obtained using funding from the Research Evaluation and Allocation Committee, the Gross Fund, and the Heart Anonymous Fund. This work was supported in part by grants from NIH (R01HL122375, R37HL078921, R01HL132832, R01HL148125) to T.M.; Uehara Memorial Foundation Fellowship and JSPS Postdoctoral Fellowship for Research Abroad to R.A.; Univ Miami funds to N.P.V.; NSF CCC (DBI-1548297), CZI BioHub Investigator Program, and Howard Hughes Medical Institute to M.P.

## Author contributions

T.M., R.A., V.N.P., and M.P. conceptualized the project. T.M. and R.A. wrote the manuscript. R.A. performed all experiments. R.A. and V.N.P. carried out the PIV analysis of cell flows. S.S. technically supported PIV analysis. V.N.P., S.S., and M.P. participated in project discussions and commented on the manuscript.

## Material and methos

### Embryo isolation and culture conditions

Fertilized eggs of White Leghorn (*Gallus Gallus domesticus*) were obtained from Petaluma Farms (Petaluma, CA) and were incubated at 37°C in a humidified incubator to the appropriate embryonic stages. We electroporated unincubated eggs at pre-streak stages IX to XII for cell flow imaging. Embryos were isolated in Tyrode’s solution (137mM NaCl, 2.7 mM KCL, 1mM MgCl2, 1.8mM CaCl2, 0.2 mM Ha2HPO4, 5.5mM D-glucose, pH 7.4). After manipulations (such as electroporation, inhibitor treatment, and implantation), the embryos were live-imaged on a vitelline membrane stretched around a glass ring according to the New culture method as previously described (33, 34).

### Implantation of COS cells

2×10^5^ of COS7 cells were plated in a 35mm cell culture dish (Thermo Fisher Scientific, 153066), 24 hours prior to transfection. The cells were transfected by Lipofectamine 3000 (Thermo Fisher Scientific, L3000015) with 6μg of pMT23-Vg1-myc-GDF1 plasmid DNA (a gift from Drs. Claudio D. Stern and Jane Dodd) for 5 hours. The next day, the transfected cells were trypsinized, and cultured using the hanging-drop method. Each hanging drop contained 10×10^6^ cells in 20 μl of culture media and were cultured overnight (O/N). The cell spheres were then rinsed in serum-free DMEM and implanted into the anterior marginal zone in the chick embryonic discs at pre-streak stages.

### Plasmid generation

ΔDEP-GFP was cloned from a plasmid, XE124 XDsh delta DEP-GFP-CS2+ in pCS2+ (a gift from Dr. Randall Moon, Addgene, #16785), by PCR (Q5 High-Fidelity DNA Polymerase, New England Biolabs, M0491) with the following primers: 5’-TACCGCGGGCCCGGGATCCAGCCACCATGGCGGAGACT-3’ and 5’-AGCCTGCACCTGAGGAGTGCTTATTTGTATAGTTCATCCATGCCATGTGTAATCC’. After PCR purification with QIAquick Gel Extraction Kit (Qiagen, #28704), the PCR product was inserted into a backbone vector, pCAG-GFP (a gift from Connie Cepko, Addgene, #1115) by using Gibson-assembly (New England Biolabs, E2611).

### Electroporation

Embryos were transfected with expression vectors, and/or control-oligo DNA conjugated with Carboxyfluorescein or Lissamine (GENE TOOLS) using an electroporator (Nepagene) with 3 pulses of 2.4-3.8 Volts, 50 milliseconds duration, 500 milliseconds interval, and platinum electrodes. The DNA solution delivered to the epiblast contained 0.1% fast green (final 0.02%), 80% glucose (final 4%) and 5μg/μl of expression vectors, or 1mM control-oligo (5’-CCTCTTACCTCAGTTACAATTTATA-3).

### Aphidicolin treatment

Aphidicolin (SIGMA, A0781), dissolved in DMSO (SIGMA, B23151), was added to Tyrode’s solution to a final concentration of 0.1 to 100μM. Embryos were isolated, and soaked in Tyrode’s solution with either 0.3% DMSO (control), or various concentrations of aphidicolin for 15 minutes at 37°C. Embryos were then cultured using the New culture method at 37°C.

### BrdU assay

Embryos were soaked in Tyrode’s solution containing BrdU (final concentration; 0.1mM, Thermo Fisher, B23151) with or without aphidicolin (SIGMA, A0781) at 37°C for 15 minutes, and cultured for 12 hours at 37°C (BrdU was incorporated for 12 hours). Embryos were then fixed in 4% paraformaldehyde (PFA, Electron Microscopy Sciences)/PBS for 30 minutes at room temperature (RT). Embryos were then washed with PBS to remove PFA, and unincorporated BrdU, and incubated with 1M HCl for 1 hour at RT to denature DNA. BrdU signal was detected by immunofluorescence staining with anti-BrdU antibody (1L200, Millipore, MAB3424).

### Tunel assay

The In Situ Cell Death Detection Kit, TMR red (Rosche, 12156792910) was used, and the kit instructions followed with these additional modifications. Embryos were fixed in 4% PFA for 30 min at RT and washed in PBS 3X for 10 minutes each. Permeabilization was performed in TBST (0.5% Triton-X) for 30 minutes at RT. Embryos were then incubated with the TUNEL reaction and DAPI staining for 3 hours at 37°C and washed with PBS five times for 10 minutes each.

### Immunofluorescent staining

#### For chicken embryos

Embryos were fixed in 4% PFA/PBS for 30 minutes at RT, or 4:1 Methanol/DMSO at 4°C O/N and washed in PBS 3X for 30 minutes each. Embryos were then incubated in blocking reagent [1% bovine serum albumin (BSA) and 0.1% triton in PBS] for 1 hour at RT. Embryos were then incubated with primary antibodies at 4°C O/N. After washing in PBS 3 X for 30 minutes each, embryos were then incubated with secondary antibodies for 2 hours at RT. Embryos were then washed in PBS 2X for 30 minutes each. For the BrdU assay, embryos had an additional incubation with Streptavidin, Alexa Fluor 647 conjugate (1:1000, Thermo Fisher Scientific) for 1 hour at RT, and washed in PBS 2X for 30 min each. Embryos were incubated with DAPI for 1 hour, washed for 30 minutes, then mounted between cover slips (VWR, 48366249), and slide glasses (Thermo Fisher Scientific, 12-544-7) with Aqua-PolyMount (Polysciences, Inc., #18606-20).

#### Primary antibodies

Anti-BrdU (1L200, Millipore, MAB3424), anti-E-Cadherin (1:500, BD Biosciences, 610181), anti-GFP (1:1000, Rockland, 600-101-215), anti-Prickle1 antibody (1:300, Proteintech, 22589-1-AP), and anti-GFP-HRP antibody (1:200, Abcam, 600-101-215), anti-ZO1 antibody (1:300, Thermo Fisher Scientific, 33-9100).

#### Secondary antibodies

Anti-mouse-IgG (H+L) Alexa Fluor 647 antibody (1:500, Thermo Fisher Scientific, A-31571), anti-goat IgG (H+L) Alexa Fluor 488 (1:500, Thermo Fisher Scientific, A11055), anti-rabbit IgG (H+L) Alexa Fluor 568 antibodies (1:500, Thermo Fisher Scientific, A10042), anti-goat IgG Alexa Fluor 488 (1:1000, Thermo Fisher Scientific, A11055) and anti-mouse IgG Alexa Fluor 594 antibodies (1:1000, Thermo Fisher Scientific, A21203).

### Whole Mount In Situ Hybridization (WISH) and subsequent immunohistochemistry

Embryos were fixed in 4% PFA/PBS in PBS O/N at 4°C. WISH was performed as previously described (39), using plasmids for *Brachyury* and *Sox3* (gift from Dr. Raymond B. Runyan), and *Vg1*. All WISH were carried out with paralleled control embryos for managing the color development time. For subsequent IHC, embryos were fixed in 4:1 methanol/DMSO at 4°C O/N.

The next day, embryos were incubated with 4:1:1 methanol/DMSO/30% Hydrogen peroxide at RT and incubated for 2 hours. Then embryos were washed in Tris-Buffered Saline (pH7.6) with 0.1% Tween 20 (TBST) 3X for 10 minutes each, and incubated with blocking solution (1% sheep serum in TBST) for 1 hour at RT. Primary antibodies, anti-GFP antibody-HRP conjugated (1:200, Abcam, ab6663), were applied for at 4°C O/N incubation. Embryos were then washed in TBST 5X for 1 hour each at RT, followed by 10 minutes with 0.05M Tris-HCl (pH 7.6), and incubated for 30 minutes with DAB (3,3′-Diaminobenzidine)/Tris-HCl (pH7.6) 0.0003% Hydrogen peroxide in the dark. After the reaction was stopped, the embryos were washed in PBS, and placed in 4% PFA at 4°C for long term storage.

### Imaging of chick embryos

For live-imaging, embryos were cultured in a 35 mm dish by the New culture method at 37°C. Time-lapse images were recorded with Nikon TIRF/Spinning Disk microscope (Nikon Ti inverted fluorescent Microscope with CSU-22 spinning disc confocal) supported by Prime 95B Scientific CMOS camera (Photometrix), Nikon Widefield Epifluorescence inverted microscope (Nikon Ti inverted fluorescent Microscope with CSU-W1 large field of view) supported by ANDOR iXon camera and Nikon Eclipse TE2000-E supported by Hamamatsu ORCA-Flash 2.8 camera. The acquisition time was every 3 minutes using Nikon Elements Advance Research software V4.00.07. All time-lapse images were recorded every 3 minutes.

Immunostained chick embryos were imaged by a confocal microscope (Leica TCS SPE). Fixed WISH samples were imaged by Leica MZ16F microscope with Leica DFC300 Fx camera and Leica FireCam V.3.4.1 software.

## Quantification and Statistical Analysis

### Cell number, mitotic rate, BrdU-, and Tunel-assay

To measure frequency of BrdU-positive or Tunel-positive cells, 3D-surface was created to an immunostained image of the embryonic disc by using the Surface tool in Imaris X64 9.2.0 software, and then the center of the embryonic disc was set for the following measurement. 3×3 squares with 800μm of each side length were then arranged on the embryonic disc. Numbers of DAPI and BrdU-positive nuclei within each square were counted by Spots tool and averaged to get a frequency of BrdU- or Tunel-positive nuclei (Fig. 2F, Fig. S1A, and Fig. S2C).

### Cell flow analyses

Cell flow analysis was performed with Imaris X64 9.2.0 software, Flowtrace and Particle Image Velocimetry (PIV) analysis. Frowtrace stacks every 20-40 frames from original live-imaging data and generates projection images of particle trajectories for visualizing cell movements. PIV analysis is described in the following part.

### PIV analysis and cell flow visualizing techniques

For quantitatively analyzing the cell flow pattern, the recorded images were further processed by using Particle Image Velocimetry (PIV) and subsequent visualization techniques. PIV analysis technique [PIVlab package in MATLAB (36)], which is a common technique for mapping displacement of particles (e.g. tagged cells) over a short time interval measured from a series of consecutive images to obtain the velocity vector field is employed to quantify the cell flows in the experimental time-lapse datasets. PIV is a standard technique used in the Fluid Physics and Engineering community to quantitatively measure and characterize fluid flows in different systems. This analysis provides comprehensive and quantitative information of a whole flow as a velocity vector field. The PIV analysis involves several steps that are carried out on image sequences over time intervals of 3 minutes, which is also the imaging acquisition speed in the experiments. First, we apply a high-pass filter during the image preprocessing step to highlight the bright particles (fluorescently tagged cell) and minimize the background. Then the PIV analysis was carried out by partitioning the image into smaller interrogation windows, typically of size 64 x 64 pixels with 50% overlap on the first pass, and a window size of 32 x 32 pixels with 50% overlap on the second pass. After this, raw velocity vector fields over time were obtained, and then these vectors were post-processed and validated by setting threshold limits, data smoothing and interpolation. A calibration was applied to convert pixel/frame units into μm/s units.

The generated velocity vector field through PIV displayed the averaged cell pattern of the whole-flow-field in the certain time period in this study. To highlight the more specific rotating cell flows, mathematical/physical information (such as topology and vorticity) was extracted from these velocity vector fields via computational visualization techniques (36, 39, 40, 42). Topology of the cell flow was determined by streamlines, which are a group of curves that approximate to contours of the vectors in the velocity vector field at a particular moment in time (39, 40, 42). Flowtrace visualized the trajectories of the cell flow, whereas the streamlines illustrated the averaged flow pattern in the time period. Therefore, the trajectories and streamlines show similar pattern when the cell flow is in stable-state; however, they are not completely consistent with each other when the cell flow is unstable and time-dependently changing (39, 40, 42). Vorticity, also termed as ‘curl of vorticity’, is a measure of a local rotation in the larger flow and its magnitude is twice that of the local angular velocity (39, 40, 42). The vorticity plot further displays directionality of the rotation. Since vorticity arises from any rotational movements, including both a full- and partial rotation, the vorticity plot displayed the vorticities in both the vortex-like rotating and curves of the cell flow (36, 39, 40, 42).

### Measurement of the distance between the left-right rotations of the polonaise movements in control-GFP and ΔDEP-embryos

The vector fields in control- and ΔDEP-GFP-embryos were generated from the recorded movies by using PIVlab and averaged the flow pattern of the polonaise movements for 6 hours. The centers of the left-right rotating cell flow of the polonaise movements in the averaged vector field were identified by streamlines and measured the distance between them by using ImageJ.

### Statistics

Statistical analyses were performed using two-tailed Student’s t tests, one-way ANOVA in R (Ver3.5.0) and Excel 2022 (Microsoft). Statistical tests, individual p values, and sample numbers are described in Figure panels and legends.

---

**Fig. S1.**
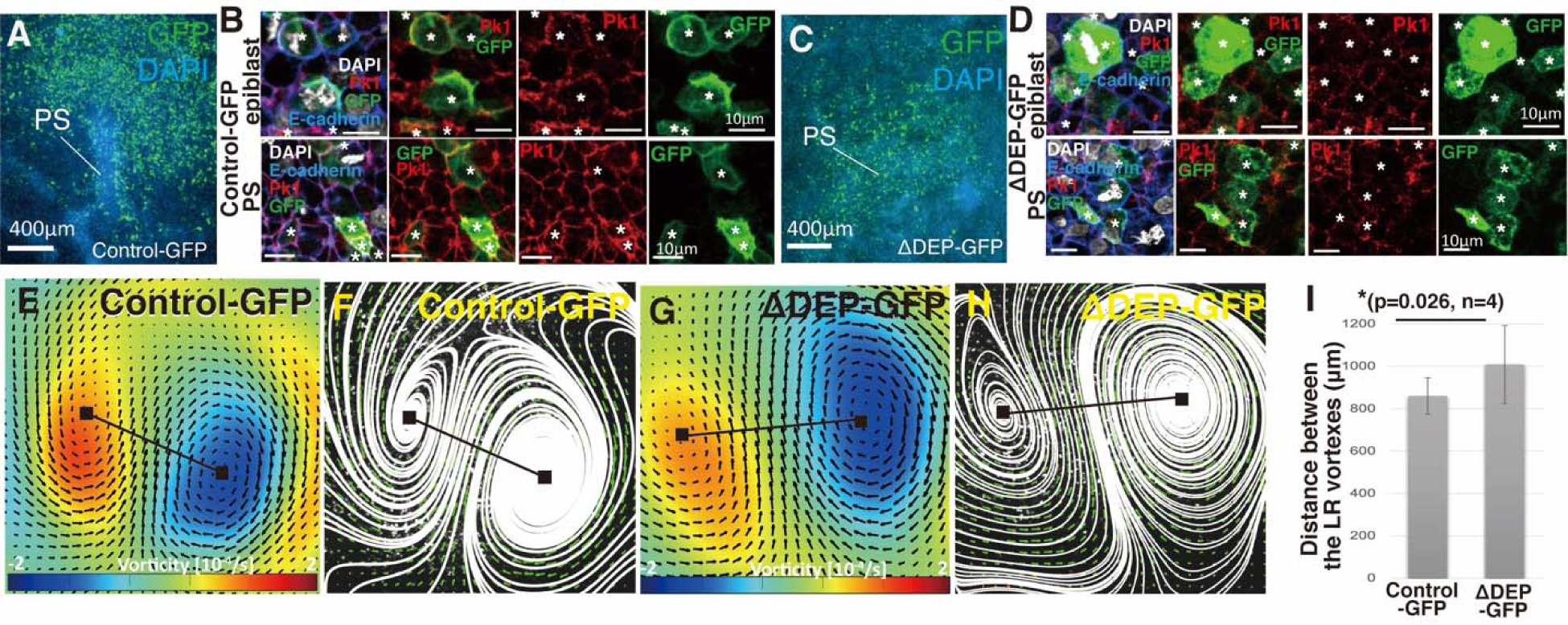
Distance between the LR rotations increases under suppression of the Wnt/PCP pathway. (A, C) Immunofluorescent staining images of control- and ΔDEP-GFP-misexpressing PSs at HH3 with GFP (green) and DAPI (cyan). (B, D) Immunostaining images of epiblast and PS cells in control- and ΔDEP-GFP-misexpressing embryos at HH3 with GFP (green), Prickle1 (Pk1; magenta), and DAPI (blue). The white asterisks indicate GFP-expressing cells. (E, G) Averaged bilateral vortex-like-rotating cell flows (6 hours), visualized by vorticity plot, in control- and ΔDEP-GFP-misexpressing embryos. Black lines, measurement of the distance between the left-right (LR) rotations. Blue, clockwise; red, counter-clockwise rotation. (F, H) Averaged bilateral vortex-like-rotating cell flows (6 hours), visualized by streamline plot, in control- and ΔDEP-GFP-misexpressing embryos. (I) Distance between the LR rotations in control- and ΔDEP-GFP-misexpressing embryos [control-GFP 860.2±86.2 μm vs ΔDEP-GFP 1008.4±183.8μm, p=0.026, n=4 for each (two-tailed Student’s t test)].

**Fig. S2.**
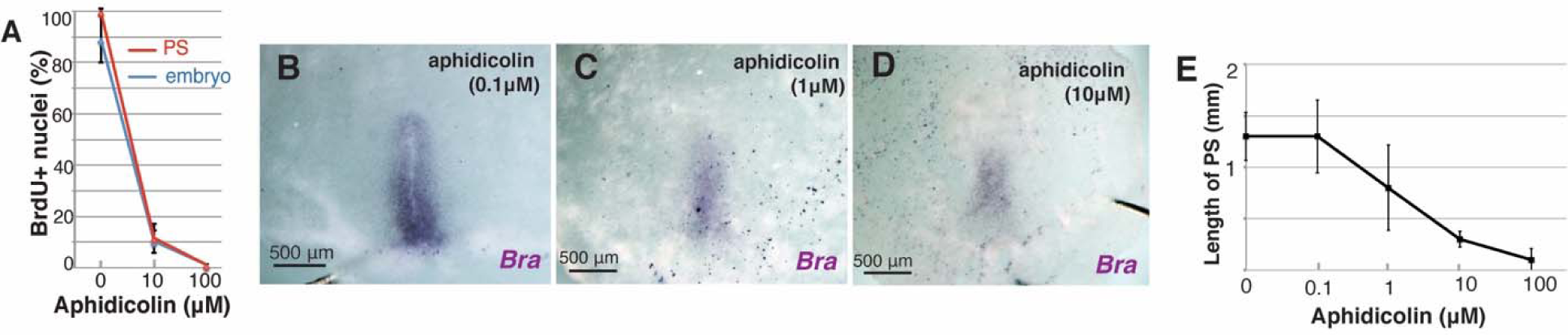
PS extension dose-dependently responds to aphidicolin. (A) Dose-dependency of aphidicolin-induced mitotic arrest (PS, control 98.8±1.3% vs 10uM aphidicolin 10.3±4.3% vs 100uM aphidicolin 0±0%, p=0.002e^-4^ in one-way ANOVA; embryo, control 87.9±9.0% vs 10uM aphidicolin 9.0±4.4% vs 100uM aphidicolin 0.02±0.01%; n=4 for each; p=0.003e^-5^ in one-way ANOVA). (B-C) In situ hybridization for Brachyury (Bra) to different concentrations of aphidicolin treatment. (D) Length of PS at HH3 for different concentrations of aphidicolin treatment (control 1.3±0.2mm vs 0.1μM aphidicolin 1.3±0.3mm vs 1μM aphidicolin 0.8±0.4mm vs 10μM aphidicolin 0.3±0.1mm vs 100μM aphidicolin 0.1±0.1mm; n=6 for each, p=6.2e^-9^ in one way ANOVA).

**Fig. S3.**
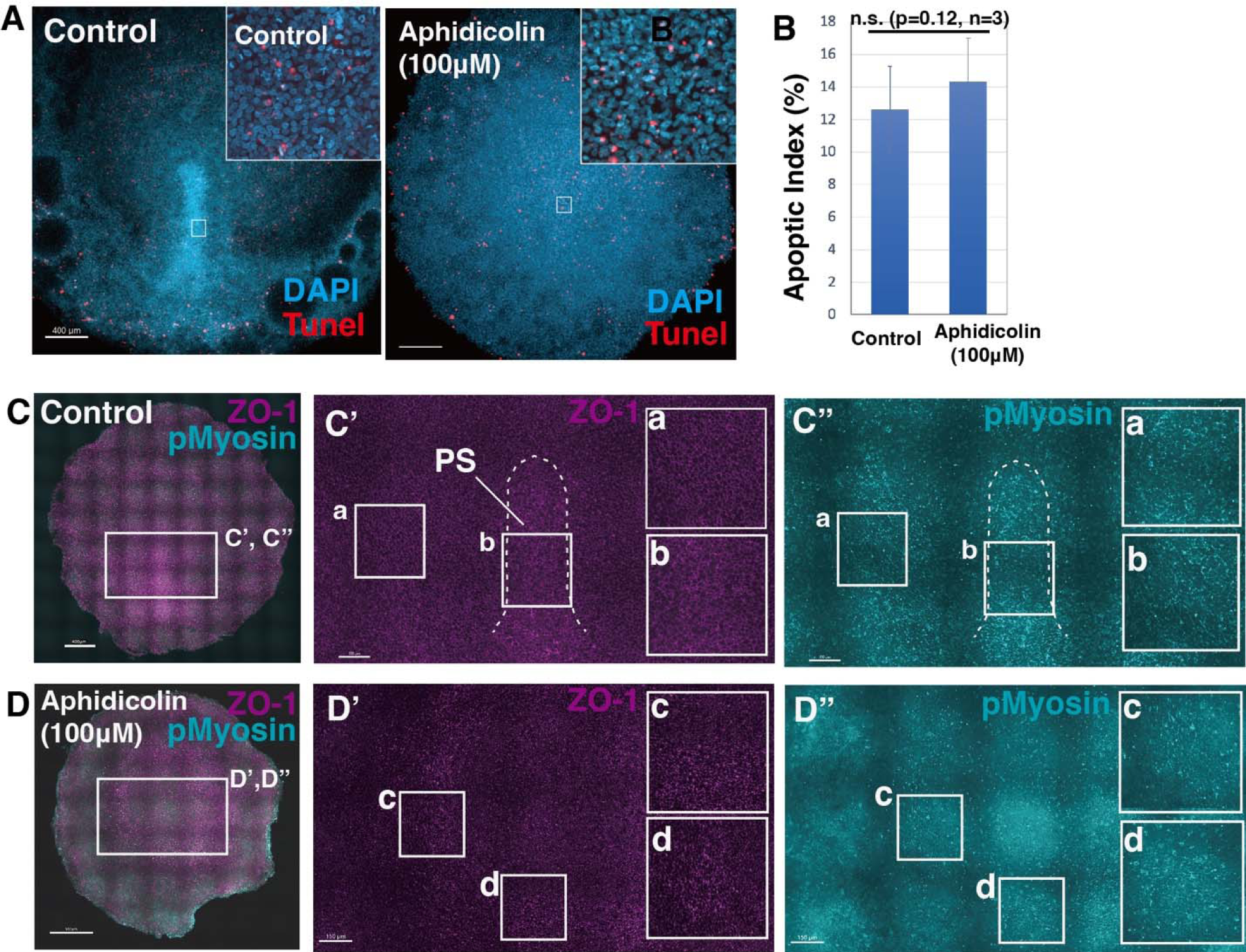
Aphidicolin-treatment maintains apoptotic index and pMyosin cables. (A) TUNEL staining at stage HH 3 in control and aphidicolin-treated embryos. (B) Apoptotic index in control and aphidicolin-treated embryos (control 10.6±3.5% vs aphidicolin 12.1±3.7%, p=0.20, n=4 for each). (C-D”) Immunofluorescent staining images of control- and aphidicolin-treated embryos at HH3 with ZO-1 (magenta) and pMyosin (cyan). White boxes show enlarged areas.

**Fig. S4.**
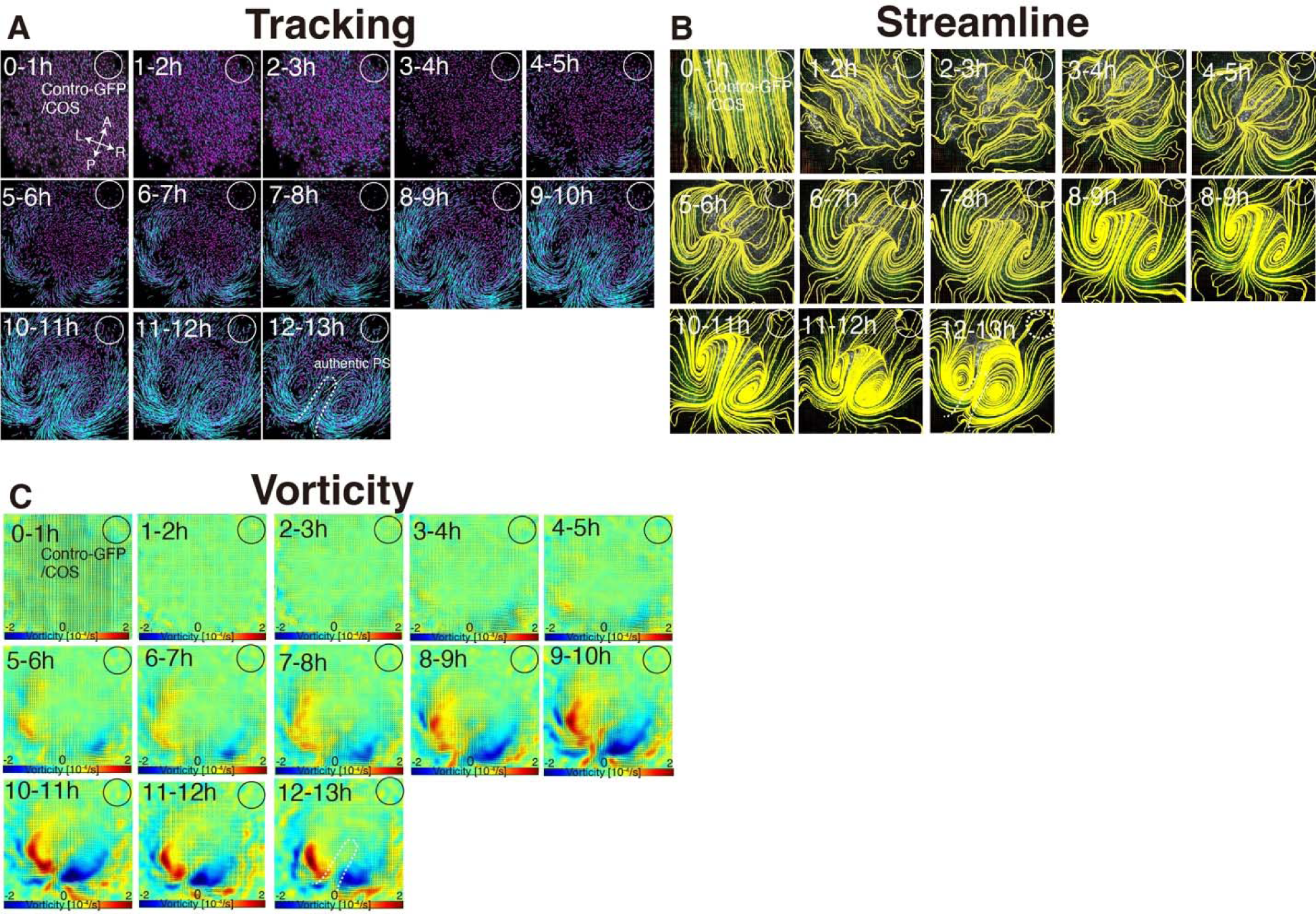
Time evolution of cell flow in control-GFP/COS-implanted embryo. (A) Trace of cell flow path of electroplated epiblast cells by using Imaris tracking analysis. See also Movie S3. A-P and L-R; anterior-posterior and left-right body axes, respectively. White circles, COS cell implanted site. (B) Streamlines, visualizing averaged cell flows during each time period. (C) Vorticity plots, displaying averaged measure of the local rotation during each time period. Blue, clockwise; red, counter-clockwise rotation.

**Fig. S5.**
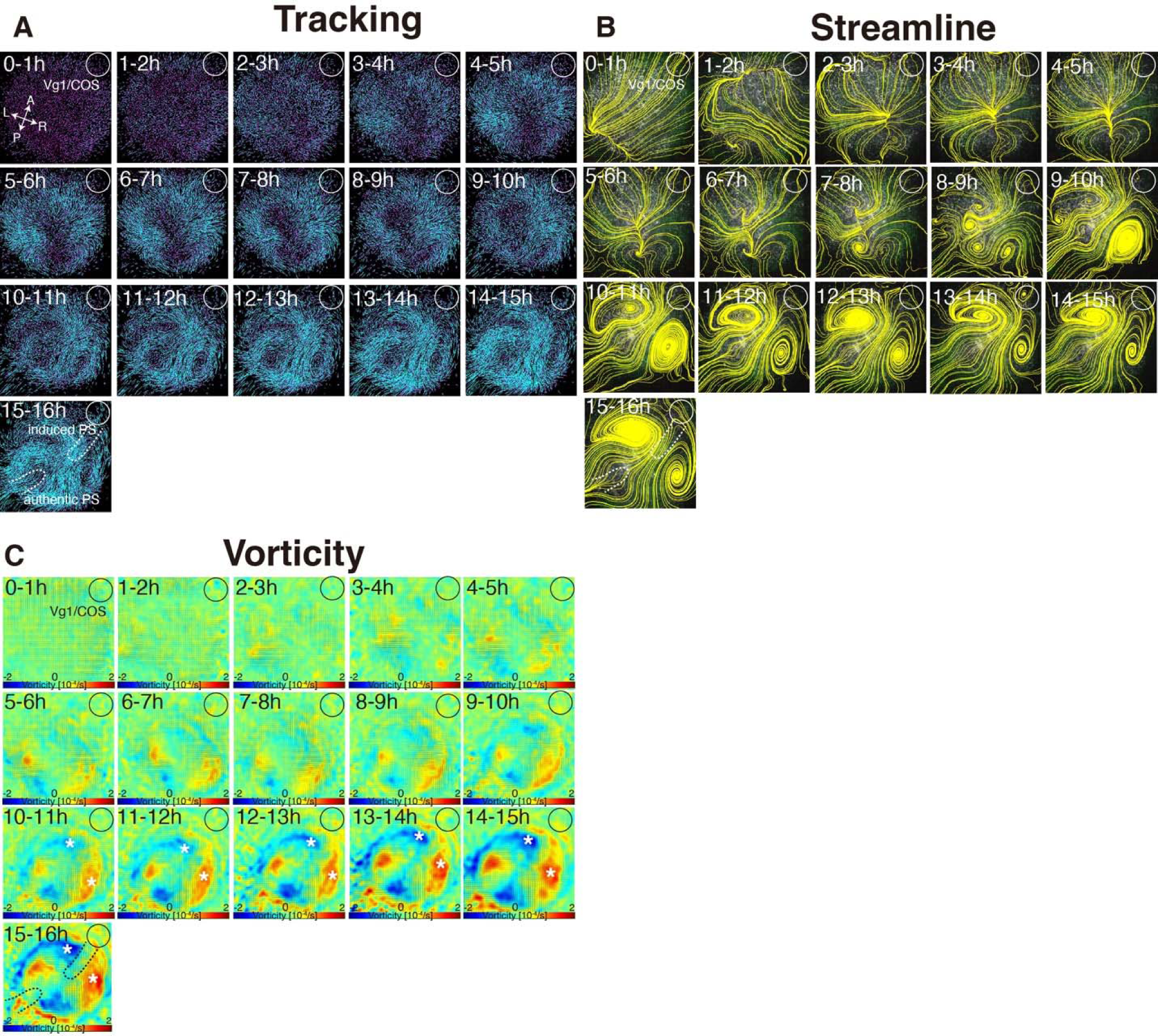
Time evolution of cell flow in Vg1/COS-implanted embryo. (A) Trace of cell flow path of electroplated epiblast cells by using Imaris tracking analysis. See also Movie S3. Axes are shown in Fig S4. White circles, COS cell implanted site. (B) Streamlines, visualizing averaged cell flows during each time period. (C) Vorticity plots, displaying averaged measure of the local rotation during each time period. Blue, clockwise; red, counter-clockwise rotation. White asterisks, the polonaise movements at the induced axis which are opposite direction to the authentic axis.

**Fig. S6.**
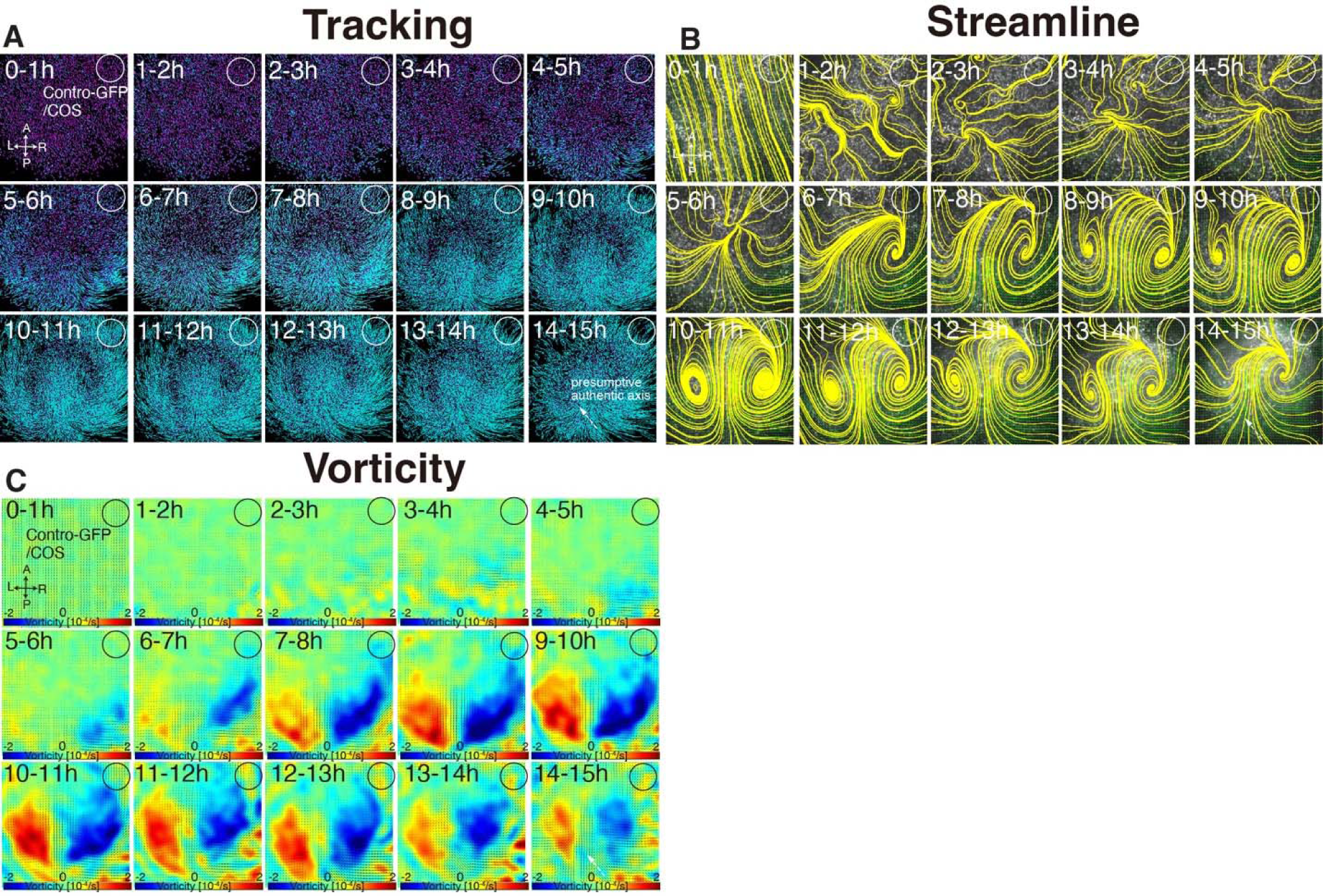
Time evolution of cell flow in control-GFP/COS-implanted embryo under aphidicolin-treatment. (A) Trace of cell flow path of electroplated epiblast cells by using Imaris tracking analysis. See also Movie S5. Axes are shown in Fig S4. White circles, COS cell implanted site. (B) Streamlines, visualizing averaged cell flows during each time period. (C) Vorticity plots, displaying averaged measure of the local rotation during each time period. Blue, clockwise; red, counter-clockwise rotation.

**Fig. S7.**
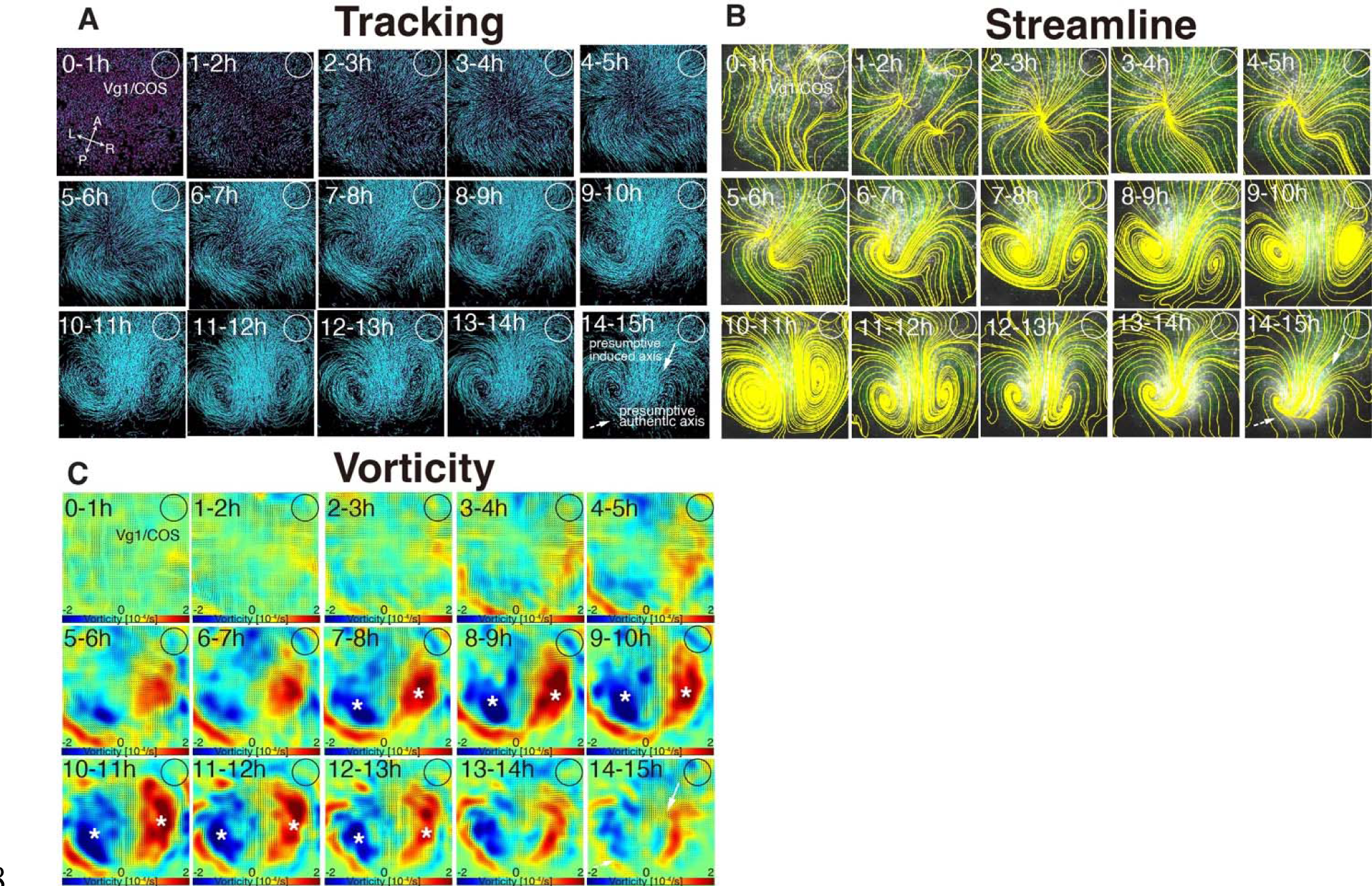
Time evolution of cell flow in Vg1/COS-implanted embryo under aphidicolin-treatment. (A) Trace of cell flow path of electroplated epiblast cells by using Imaris tracking analysis. See also Movie S5. Axes are shown in Fig S4. White circles, COS cell implanted site. (B) Streamlines, visualizing averaged cell flows during each time period. (C) Vorticity plots, displaying averaged measure of the local rotation during each time period. Blue, clockwise; red, counter-clockwise rotation. White asterisks, the polonaise movements at the induced axis which are opposite direction to the authentic axis.

## Supplemental movies

**Movie S1. ‘Polonaise movements’ continue under suppression of Wnt/PCP pathway.** Time-lapse movie was analyzed by using Imaris tracking. Embryos were co-electroporated with lissamine-tagged oligo (red) and control-GFP or ΔDEP-GFP expression vectors (cyan) at pre-streak stage X-XII. Live imaging performed at 4x objective lens on a spinning disk confocal microscope until HH3. Related to Fig. 1 and Fig. S1. Scale bars, 300 μm.

**Movie S2. Mitotic arrest maintains early phase of ‘polonaise movements’**. Time-lapse movie of fluorescently-tagged epiblast cells in control and aphidicolin-treated embryos from pre-streak to HH3, processed by using Flowtrace. The trajectories, from green to yellow, indicate movement of fluorescently tagged cells (30 min projections). Embryos were electroporated with fluorescein-tagged oligo (green) and treated with DMSO (control-sham operated) or 100 μM of aphidicolin. Live-imaging performed at 4x objective lends on a spinning disk confocal microscope for 10 hours. Related to Fig. 2, Fig. S2, and S3. Scale bars, 200 μm.

**Movie S3. Authentic PS extends in absence of regular ‘polonaise movements’.** Embryos were electroporated with lissamine-tagged oligo (red) and implanted with COS cells expressing either a control-GFP (control-GFP/COS) or Vg1 (Vg1/COS) expression vector at anterior marginal zone at pre-streak stage IX-X. Live-imaging performed at a 2x objective lens on a wide-field epifluorescent microscope. Magenta dots; lissamine-tagged epiblast cells, cyan lines; trajectories of the tagged epiblast cells for 2 hours. Related to Fig. 3, Fig. S4, S5. Scale bars, 300 μm.

**Movie S4. Ectopic Vg1 leads to ‘polonaise movements’ despite defective PS morphogenesis after mitotic arrest.** Embryos were electroporated with lissamine-tagged oligo (red), implanted with either control-GFP/COS or Vg1/COS at the anterior marginal zone (AMZ), and treated with 100 μM of aphidicolin at pre-streak stage IX to X. Live imaging performed at a 2x objective lens with a wide-field epifluorescent microscope. Magenta dots; lissamine tagged epiblast cells, cyan lines; trajectories for 2.5 hours. Related to Fig. 4, Fig. S6, and S7. Scale bars, 300 μm.

## References

1. S. F. Gilbert, M. J. F. Barresi, Developmental Biology (Oxford University Press, 2017).

2. L. Solnica-Krezel, D. S. Sepich, Gastrulation: making and shaping germ layers. Annu Rev Cell Dev Biol 28, 687–717 (2012).

3. M. Leptin, Gastrulation movements: the logic and the nuts and bolts. Dev Cell 8, 305–320 (2005).

4. L. Solnica-Krezel, Conserved patterns of cell movements during vertebrate gastrulation. Curr Biol 15, R213–228 (2005).

5. R. Keller, L. Davidson, Cell movements of gastrulation. Gastrulation: from cells to embryos (ed. C. Stern), 291-304 (2004).

6. T. Mikawa, A. M. Poh, K. A. Kelly, Y. Ishii, D. E. Reese, Induction and patterning of the primitive streak, an organizing center of gastrulation in the amniote. Dev Dyn 229, 422–432 (2004).

7. R. Keller, Shaping the Vertebrate Body Plan by Polarized Embryonic Cell Movements. Science 298, 1950–1954 (2002).

8. E. W. Gehrels, B. Chakrabortty, M. E. Perrin, M. Merkel, T. Lecuit, Curvature gradient drives polarized tissue flow in the Drosophila embryo. Proc Natl Acad Sci U S A 120, e2214205120 (2023).

9. C. Bertet, L. Sulak, T. Lecuit, Myosin-dependent junction remodelling controls planar cell intercalation and axis elongation. Nature 429, 667–671 (2004).

10. R. Keller et al., Mechanisms of convergence and extension by cell intercalation. Philos Trans R Soc Lond B Biol Sci 355, 897–922 (2000).

11. M. Tada, M. L. Concha, C.-P. Heisenberg (2002) Non-canonical Wnt signalling and regulation of gastrulation movements. in Seminars in cell & developmental biology (Elsevier), pp 251-260.

12. R. Wetzel, Untersuchungen am Hühnchen. Die Entwicklung des Keims während der ersten beiden Bruttage. Wilhelm Roux Arch Entwickl Mech Org 119, 188–321 (1929).

13. L. Gräper, Die Primitiventwicklung des Hühnchens nach stereokinematographischen Untersuchungen, kontrolliert durch vitale Farbmarkierung und verglichen mit der Entwicklung anderer Wirbeltiere. Wilhelm Roux Arch Entwickl Mech Org 116, 382–429 (1929).

14. C. D. Stern, Gastrulation in the chick. C. D. Stern, Ed., Gastrulation: from cells to embryo (CSHL Press, New York, 2004), pp. 219–232.

15. G. C. Schoenwolf, Molecular Genetic Control of Axis Patterning during Early Embryogenesis of Vertebrates. Annals of the New York Academy of Sciences 919, 246–260 (2000).

16. A. Raffaelli, C. D. Stern, Signaling events regulating embryonic polarity and formation of the primitive streak in the chick embryo. Curr Top Dev Biol 136, 85–111 (2020).

17. G. Sheng, A. Martinez Arias, A. Sutherland, The primitive streak and cellular principles of building an amniote body through gastrulation. Science 374, abg1727 (2021).

18. E. J. Sanders, M. Varedi, A. S. French, Cell proliferation in the gastrulating chick embryo: a study using BrdU incorporation and PCNA localization. Development 118, 389–399 (1993).

19. C. Cui, X. Yang, M. Chuai, J. A. Glazier, C. J. Weijer, Analysis of tissue flow patterns during primitive streak formation in the chick embryo. Dev Biol 284, 37–47 (2005).

20. M. Saadaoui, D. Rocancourt, J. Roussel, F. Corson, J. Gros, A tensile ring drives tissue flows to shape the gastrulating amniote embryo. Science 367, 453–458 (2020).

21. O. Voiculescu, F. Bertocchini, L. Wolpert, R. E. Keller, C. D. Stern, The amniote primitive streak is defined by epithelial cell intercalation before gastrulation. Nature 449, 1049–1052 (2007).

22. R. O’Rahilly, F. Muller, Developmental stages in human embryos: revised and new measurements. Cells Tissues Organs 192, 73–84 (2010).

23. R. Asai, M. Bressan, T. Mikawa, Avians as a Model System of Vascular Development. Methods Mol Biol 2206, 103–127 (2021).

24. V. Hamburger, H. L. Hamilton, A series of normal stages in the development of the chick embryo. 1951. Dev Dyn 195, 231–272 (1992).

25. E. A. Seleiro, D. J. Connolly, J. Cooke, Early developmental expression and experimental axis determination by the chicken Vg1 gene. Curr Biol 6, 1476–1486 (1996).

26. A. Streit et al., Chordin regulates primitive streak development and the stability of induced neural cells, but is not sufficient for neural induction in the chick embryo. Development 125, 507–519 (1998).

27. M. Chuai, G. Serrano Nájera, M. Serra, L. Mahadevan, C. J. Weijer, Reconstruction of distinct vertebrate gastrulation modes via modulation of key cell behaviors in the chick embryo. Sci Adv 9, eabn5429 (2023).

28. E. Rozbicki et al., Myosin-II-mediated cell shape changes and cell intercalation contribute to primitive streak formation. Nat Cell Biol 17, 397–408 (2015).

29. O. Voiculescu, L. Bodenstein, I. J. Lau, C. D. Stern, Local cell interactions and self-amplifying individual cell ingression drive amniote gastrulation. Elife 3, e01817 (2014).

30. O. Voiculescu, Movements of chick gastrulation. Curr Top Dev Biol 136, 409–428 (2020).

31. J. X. Wang, M. D. White, Mechanical forces in avian embryo development. Semin Cell Dev Biol 120, 133–146 (2021).

32. H. Eyal-Giladi, S. Kochav, From cleavage to primitive streak formation: a complementary normal table and a new look at the first stages of the development of the chick. I. General morphology. Dev Biol 49, 321–337 (1976).

33. D. A. T. New, A New Technique for the Cultivation of the Chick Embryo in vitro. Journal of Embryology and Experimental Morphology 3, 326–331 (1955).

34. L. Maya-Ramos, T. Mikawa, Programmed cell death along the midline axis patterns ipsilaterality in gastrulation. Science 367, 197–200 (2020).

35. W. Gilpin, V. N. Prakash, M. Prakash, Flowtrace: simple visualization of coherent structures in biological fluid flows. J Exp Biol 220, 3411–3418 (2017).

36. W. Thielicke, E. J. Stamhuis, PIVlab – Towards User-friendly, Affordable and Accurate Digital Particle Image Velocimetry in MATLAB. Journal of Open Research Software 2 (2014).

37. E. J. Stamhuis, Basics and principles of particle image velocimetry (PIV) for mapping biogenic and biologically relevant flows. Aquatic Ecology 40, 463–479 (2006).

38. M. Raffel, C. E. Willert, J. Kompenhans, Particle image velocimetry: a practical guide (Springer, 1998), vol. 2.

39. C. K. Batchelor, G. K. Batchelor, An introduction to fluid dynamics 2nd edition (Cambridge university press, 2012).

40. 40. R. Aris, Vectors, tensors and the basic equations of fluid mechanics (Courier Corporation, 2012).

41. M. Jiang, R. Machiraju, D. Thompson, Detection and visualization of Vortices. The visualization handbook 295 (2005).

42. H. G. W. Christopher J. Greenshields, Notes on Computational Fluid Dynamics: General Principles (CFD Direct Limited, 2022).

43. M. Sharma, I. Castro-Piedras, G. E. Simmons, Jr., K. Pruitt, Dishevelled: A masterful conductor of complex Wnt signals. Cell Signal 47, 52–64 (2018).

44. U. Rothbächer et al., Dishevelled phosphorylation, subcellular localization and multimerization regulate its role in early embryogenesis. Embo j 19, 1010–1022 (2000).

45. Y. Yang, M. Mlodzik, Wnt-Frizzled/Planar Cell Polarity Signaling: Cellular Orientation by Facing the Wind (Wnt). Annual Review of Cell and Developmental Biology 31, 623–646 (2015).

46. J. M. Glynn, T. G. Cotter, D. R. Green (1992) Apoptosis induced by actinomycin D, camptothecin or aphidicolin can occur in all phases of the cell cycle. (Portland Press Ltd.).

47. J. Cooke, Morphogenesis and regulation in spite of continued mitotic inhibition in Xenopus embryos. Nature 242, 55–57 (1973).

48. Y. Wei, T. Mikawa, Formation of the avian primitive streak from spatially restricted blastoderm: evidence for polarized cell division in the elongating streak. Development 127, 87–96 (2000).

49. S. B. Shah et al., Misexpression of chick Vg1 in the marginal zone induces primitive streak formation. Development 124, 5127–5138 (1997).

50. G. S. Nájera, C. J. Weijer, The evolution of gastrulation morphologies. Development 150 (2023).

51. I. Roszko, A. Sawada, L. Solnica-Krezel, Regulation of convergence and extension movements during vertebrate gastrulation by the Wnt/PCP pathway. Semin Cell Dev Biol 20, 986–997 (2009).

52. M. Chuai et al., Cell movement during chick primitive streak formation. Dev Biol 296, 137–149 (2006).

53. Y. Ishii, T. Mikawa, Somatic transgenesis in the avian model system. Birth Defects Res C Embryo Today 75, 19–27 (2005).

54. C. F. Davey, C. B. Moens, Planar cell polarity in moving cells: think globally, act locally. Development 144, 187–200 (2017).

55. M. Tada, C.-P. Heisenberg, Convergent extension: using collective cell migration and cell intercalation to shape embryos. Development 139, 3897–3904 (2012).

56. M. A. Hartman, J. A. Spudich, The myosin superfamily at a glance. Journal of Cell Science 125, 1627–1632 (2012).

57. P. Caldarelli, A. Chamolly, O. Alegria-Prévot, J. Gros, F. Corson, Self-organized tissue mechanics underlie embryonic regulation. bioRxiv 10.1101/2021.10.08.463661, 2021.2010.2008.463661 (2021).

58. J. Green, Morphogen gradients, positional information, and Xenopus: Interplay of theory and experiment. Developmental Dynamics 225, 392–408 (2002).

## References for Material and methods

1. S. F. Gilbert, M. J. F. Barresi, Developmental Biology (Oxford University Press, 2017).

3. M. Leptin, Gastrulation movements: the logic and the nuts and bolts. Dev Cell 8, 305–320 (2005).

63. R. Keller, L. Davidson, Cell movements of gastrulation. Gastrulation: from cells to embryos (ed. C. Stern), 291-304 (2004).

69. M. Tada, M. L. Concha, C.-P. Heisenberg (2002) Non-canonical Wnt signalling and regulation of gastrulation movements. in Seminars in cell & developmental biology (Elsevier), pp 251-260.

72. C. D. Stern, Gastrulation in the chick. C. D. Stern, Ed., Gastrulation: from cells to embryo (CSHL Press, New York, 2004), pp. 219–232.

30. O. Voiculescu, Movements of chick gastrulation. Curr Top Dev Biol 136, 409–428 (2020).

96. M. Raffel, C. E. Willert, J. Kompenhans, Particle image velocimetry: a practical guide (Springer, 1998), vol. 2.

98. R. Aris, Vectors, tensors and the basic equations of fluid mechanics (Courier Corporation, 2012).

104. J. M. Glynn, T. G. Cotter, D. R. Green (1992) Apoptosis induced by actinomycin D, camptothecin or aphidicolin can occur in all phases of the cell cycle. (Portland Press Ltd.).

50. G. S. Nájera, C. J. Weijer, The evolution of gastrulation morphologies. Development 150 (2023).

